# Salivary proteins in spider mites: dual roles in feeding and silk fiber coating

**DOI:** 10.1101/2025.02.07.637200

**Authors:** Yuka Arai, Naoki Takeda, Shinya Komoto, Hisashi Murakami, Dagmar Voigt, Takeshi Suzuki

**Affiliations:** Graduate School of Bio-Applications and Systems Engineering, Tokyo University of Agriculture and Technology, Koganei, Tokyo, Japan; Imaging Core Facility, Okinawa Institute of Science and Technology Graduate University (OIST), Tancha, Onna-son, Okinawa, Japan; Botany, Faculty of Biology, Technische Universität Dresden, Dresden, Germany; Institute of Global Innovation Research, Tokyo University of Agriculture and Technology, Koganei, Tokyo, Japan

**Keywords:** atomic force microscope (AFM), cryo-SEM, Fibroin, *in situ* hybridization, micro-CT, plant cell-sucking, proteomics, RNAi, saliva, spider mite

## Abstract

Spider mites are plant cell-sucking herbivores that spin nanoscale adhesive silk fibers released from their mouthpart appendages. Previous whole-genome sequencing predicted 17 silk genes in the two-spotted spider mite, *Tetranychus urticae*, while silk proteomics detected two candidates, Fibroin-1 and Fibroin-2. However, saliva proteomics also detected Fibroin-1, Fibroin-2, and sFibroin-1 (with high sequence similarity to Fibroin-1). We investigated whether these three proteins function in silk, saliva, or both. We found *Fibroin-1*, *sFibroin-1*, and *Fibroin-2* genes expressed in the salivary glands but found no direct internal connection between salivary and silk glands. Our silk proteomics detected 8 silk proteins previously predicted and 66 salivary proteins, including Fibroin-1, sFibroin-1, and Fibroin-2. We detected a bead-on-a-string pattern formed by fluid (hypothesized to be saliva) on the silk fiber and a fluid patch that remains at the piercing site on the leaf surface after mite feeding. RNAi-mediated silencing of *Fibroin-1* and *sFibroin-1* genes reduced mite feeding duration, survival, and fecundity, and silencing of *Fibroin-2* gene showed a tendency to reduce the thickness of individual silk fibers. These results suggest that Fibroin-1, sFibroin-1, and Fibroin-2 are secreted from the salivary glands and then adhere to silk fibers. Fibroin-1 and sFibroin-1 may assist mites to adhere the tip of the mouthparts to the leaf surface when sucking with their stylets, while Fibroin-2 may coat silk fibers. This study reveals the dual role of saliva in plant cell-sucking herbivorous mites, namely, stabilizing the mouthparts during feeding and coating silk fibers to provide them with adhesive properties.

**Highlights:** Three salivary proteins of spider mites are abundantly detected in their silk fibers.

Though the salivary glands and silk glands are not connected, saliva appears to adhere to silk fibers after secretion.

Knockdown of two of the three salivary protein genes suppressed the feeding behavior of spider mites, resulting in reduced survival and fecundity.

Knockdown of the remaining salivary protein gene showed a tendency to reduce the thickness of the silk filament.

## Introduction

Silk is a fibrous protein that plays a crucial role in the survival and reproduction of hundreds of thousands of arthropods.^1,2^ Silk proteins contain highly repetitive sequences of amino acids, are stored as liquid in silk glands, and are configured into fibers when spun upon secretion.^3^ The genes encoding silk proteins have evolved independently numerous times throughout arthropod evolution. In silkworms, fibroin heavy chain, a fibrous protein of silk, contains repetitive motifs such as Gly-Ala-Gly-Ala-Gly-Ala-Ser and forms β-sheet crystals,^4,5^. In contrast, spidroin, a fibrous protein of spiders, has β-sheet structures with poly-Ala and poly-Gly-Ala motifs and helical structures with repetitive motifs such as Gly-Gly-X (where X is Ala, Asp, or Tyr).^4,6^

Spider mites (Trombidiformes, Tetranychidae), herbivorous chelicerates, feed on the contents of plant cells using a pair of first appendages called chelicerae that form a stylet and extend from the mouthparts.^47^ Spider mites spin adhesive silk for locomotion,^7^ dispersal,^8,9^ attracting males,^10^ capturing and eliminating competitors,^11,12^ and preventing predator invasion.^13–16^ The spider mite silk gland is a large single cell containing electron-lucent grana, rough endoplasmic reticulum, and mitochondria. The duct leading from the silk gland opens at the tip of the spinnerets, which are located in a pair of second appendages called pedipalps that extend from the mouthparts.^17^ Spider mites continuously spin nanoscale silk fibers from the spinnerets as they walk.^7,18,19^

In the fully sequenced and annotated genome of the two-spotted spider mite (TSSM) *Tetranychus urticae* Koch, 17 silk protein genes were computationally predicted.^20^ The amino acid sequences of the 17 genes contain repetitive motifs that are rich in Gly, Ala, and Ser, similar to those in the silkworm fibrous silk protein. However, silk proteomics has identified two candidate silk proteins, Fibroin-1 and Fibroin-2, that contain repetitive motifs rich in Val, Asn, and Ser and are distinct from any of the 17 originally predicted genes.^21^ Furthermore, these two proteins and a protein exhibiting 72.1% similarity to Fibroin-1, called sFibroin-1, were detected in the TSSM saliva proteome.^22,23^ Current knowledge based on these findings is that Fibroin-1, sFibroin-1, and Fibroin-2 are secreted as saliva and that at least Fibroin-1 and Fibroin-2 are also present in the silk fibers of TSSM. This background raises the following questions about TSSM saliva and silk: 1) What are the functions of Fibroin-1, sFibroin-1, and Fibroin-2 in the saliva and/or the silk? 2) How are these proteins incorporated into silk fibers? 3) Which silk proteins are produced in the silk gland and secreted through the spinnerets?

Our investigation involves four steps to address these questions. First, we visualized the nanoscale silk fibers of TSSM with cryo-scanning electron microscopy (cryo-SEM) and measured their dimensions with atomic force microscopy (AFM). In addition, we performed micro-computed tomography (CT) analysis to search for morphological evidence of connections between the salivary gland and the silk gland. Second, we conducted spatial analysis of *Fibroin-1*, *sFibroin-1*, and *Fibroin-2* gene expression by *in situ* hybridization (ISH). Third, we performed silk proteomics to quantify the silk proteins and salivary proteins adhered to silk fibers. Fourth, we performed RNA interference (RNAi)-mediated functional analysis of *Fibroin-1*, *sFibroin-1*, and *Fibroin-2* genes in the context of mite feeding and the interaction with silk spinning.

## Results

### Morphological observations of silk fibers and saliva

TSSM releases silk fibers from spinnerets located on the pedipalps, a pair of second appendages in the mouthparts (Fig. 1A). Cryo-SEM imaging revealed silk fibers composed of single filaments and bundles of varying numbers of filaments (Fig. 1B). AFM imaging showed that the average diameter of a single silk filament spun by adult females was 61.9 ± 1.8 nm (Fig. 1C). Cryo-SEM imaging visualized a bead-on-a-string pattern formed by drops of fluid on several silk fibers (Fig. 1D), as well as fluid patches remaining at silk fixation sites on challenging substrates, such as tomato trichomes (Fig. 1E), and at the piercing site established for mite feeding (Fig. 1F, G). We hypothesized that the fluid was saliva.

**Figure 1.**
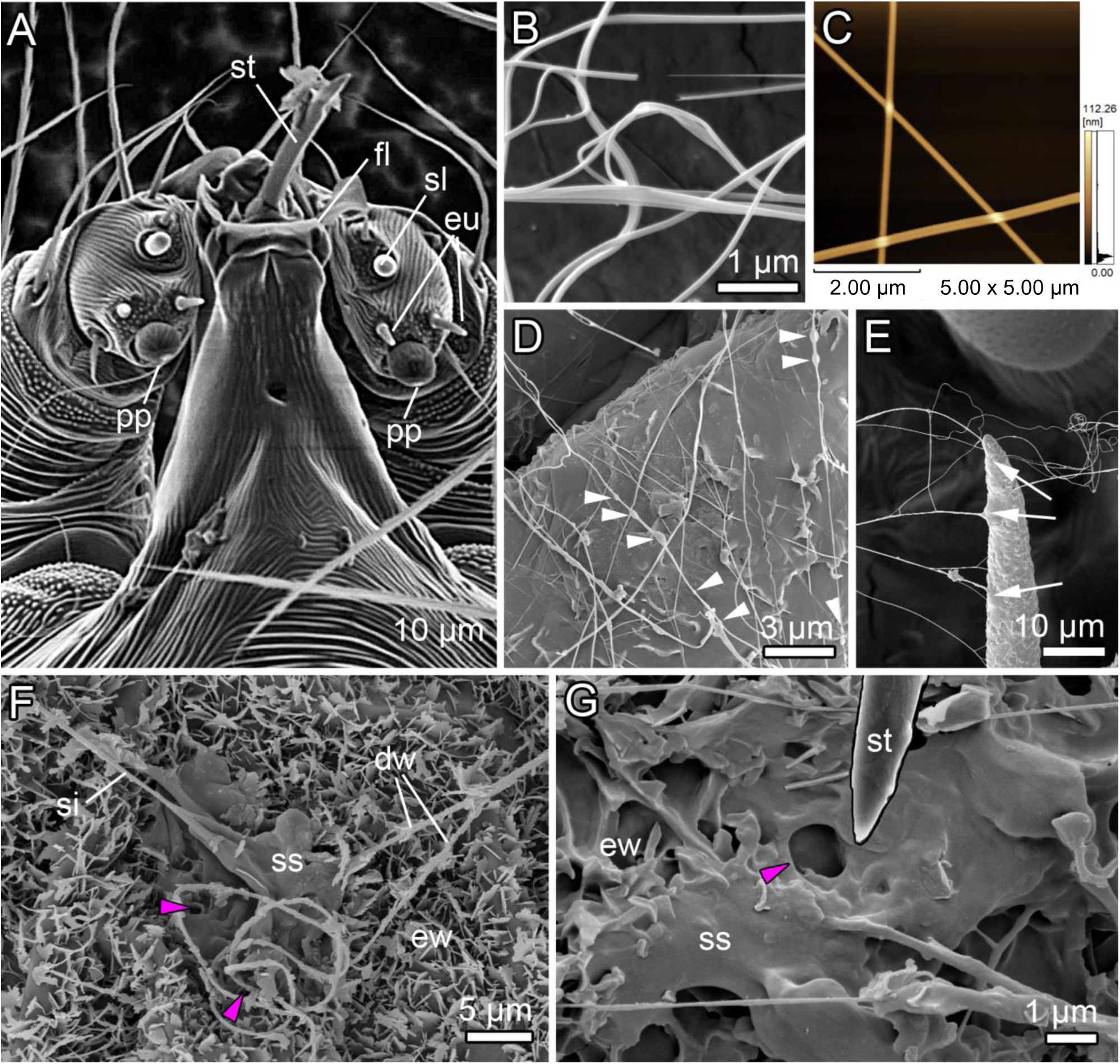
Silk and saliva morphology. (A) Scanning electron microscopy (SEM) image of the ventral side of a TSSM gnathosoma. TSSM releases silk fibers from a pair of spinnerets (arrows) at the tips of the pedipalps (pp), the second appendage. Also visible are the eupathidium (eu), flap (fl), solenidion (sl), and stylets (st). (B) Cryo-SEM image of adult TSSM silk fibers, showing single filaments and bundles with varying numbers of filaments. (C) Atomic force microscopy image of adult TSSM silk fibers. The average diameter of a single filament was 61.9 ± 1.8 nm (*n* = 9). (D) Cryo-SEM image of adult TSSM silk fibers on the adaxial surface of a beach strawberry leaf. Fibers are in contact with the convex epidermal cell parts, which are covered with crystalline wax ribbons. White arrowheads indicate fluid on the silk fibers, forming a beads-on-a-string pattern. (E) Cryo-SEM image of adult TSSM silk fibers and bundles attached to a non-glandular tomato trichome tip (ct). Arrows mark fluid patches from which silk fibers arise. (F, G) Cryo-SEM images of adult TSSM silk fibers (si) crossing the fluid patch formed around a stylet insertion site (magenta arrowheads) on the abaxial surface of a beach strawberry leaf covered with ribbon-shaped epicuticular wax crystals (ew). Note the detached wax crystals (dw) adhering to the fluid on the silk fibers.G shows a detail of F with the tentatively identified silk-saliva mixture (ss) and the stylet tip of the TSSM (st).

### *ISH of* Fibroin-1, sFibroin-1, *and* Fibroin-2 *genes and the internal morphology of the proterosoma*

ISH showed that *Fibroin-1*, *sFibroin-1*, and *Fibroin-2* genes were expressed in the dorsal podocephalic glands, a pair of salivary glands (Fig. 2A-J). Micro-CT scanning visualized the stylet (Fig. 2K) and salivary ducts (Fig. 2L). The stylet originates from the stylophore, runs posteriorly, bends approximately 180°, and then runs anteriorly to the anterior end of the rostrum. The salivary duct arises from the dorsal and anterior podocephalic glands, runs downward, bends approximately 180°, and then runs upward parallel to the stylet to the anterior end of the rostrum. No direct connection between the salivary glands and the silk glands was observed, so saliva and silk meet each other during simultaneous secretion and spinning.

**Figure 2.**
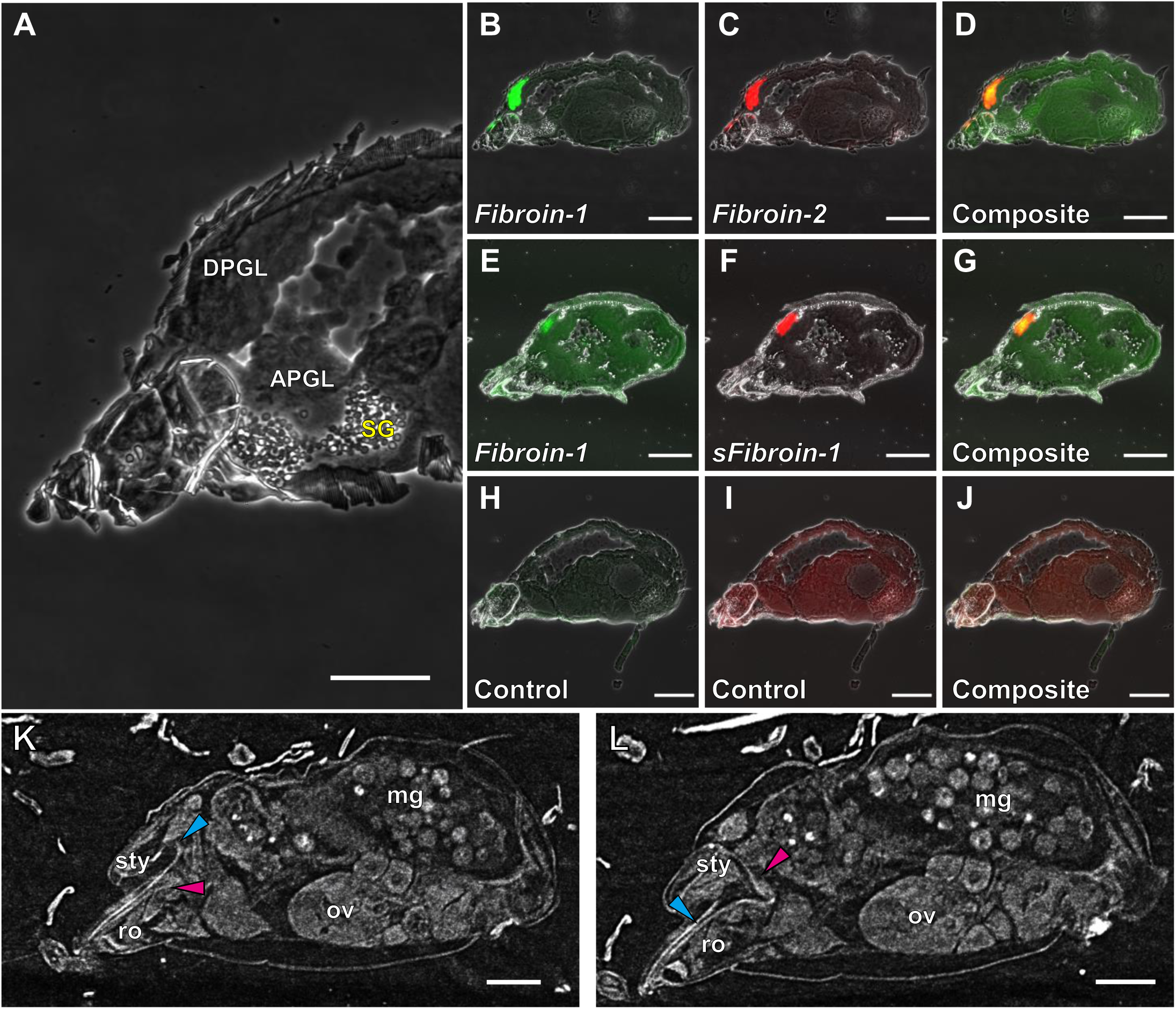
Internal morphology of the proterosoma and localization of *Fibroin-1*, *sFibroin-1*, and *Fibroin-2* gene expression. (A) A sagittal cross-section of the proterosoma of an adult female TSSM shows the dorsal podocephalic gland (DPGL), anterior podocephalic gland (APGL), and silk gland (SG). The cytoplasm of SG contains many secretory vesicles filled with liquid silk. (B–J) RNAscope *in situ* hybridization shows the localization of *Fibroin-1*, *sFibroin-1*, and *Fibroin-2* gene expression and a negative control. Superimposed (composite) images (D, G, and J) show co-staining in the same cross-section. In B, E, and H, fluorescence images at 519 nm were obtained at an excitation wavelength of 495 nm. In C, F, and I, fluorescence images at 578 nm were obtained at an excitation wavelength of 552 nm. (K, L) Sagittal cross-sectional micro-computed tomography images taken at different angles in an adult TSSM female show the stylet (cyan arrowhead) and the salivary duct (magenta arrowhead). In K, the stylet originates at the stylophore (sty), runs backwards, then bends about 180° to run forwards to the front end of the rostrum (ro). In L, the salivary duct originates from the podocephalic gland, runs downwards, then bends about 180° to run upwards, running parallel to the stylet to the anterior end of the rostrum. The midgut (mg) contains multiple large floating cells called digestive cells. ov, ovary. Scale bars: (A, K, L) 50 μm; (B–J) 100 μm.

### Silk proteomics

The salivary proteins Fibroin-1, sFibroin-1, and Fibroin-2 were detected in the proteome of silk fibers (Fig. 3). The abundance of each was higher in samples of silk fibers than in samples taken from the whole body. Eight of the 17 candidate silk proteins^20^ were detected in the proteome of silk fibers, and six of the eight proteins were more abundant in silk fibers than in whole-body samples. However, the abundance of each of the eight proteins was lower than that of Fibroin-1, sFibroin-1, or Fibroin-2 in both silk fibers and whole-body samples. In addition to Fibroin-1, sFibroin-1, and Fibroin-2, our silk proteome included 66 of the 95 salivary proteins identified by saliva proteomics^22^ (Fig. S1).

**Figure 3.**
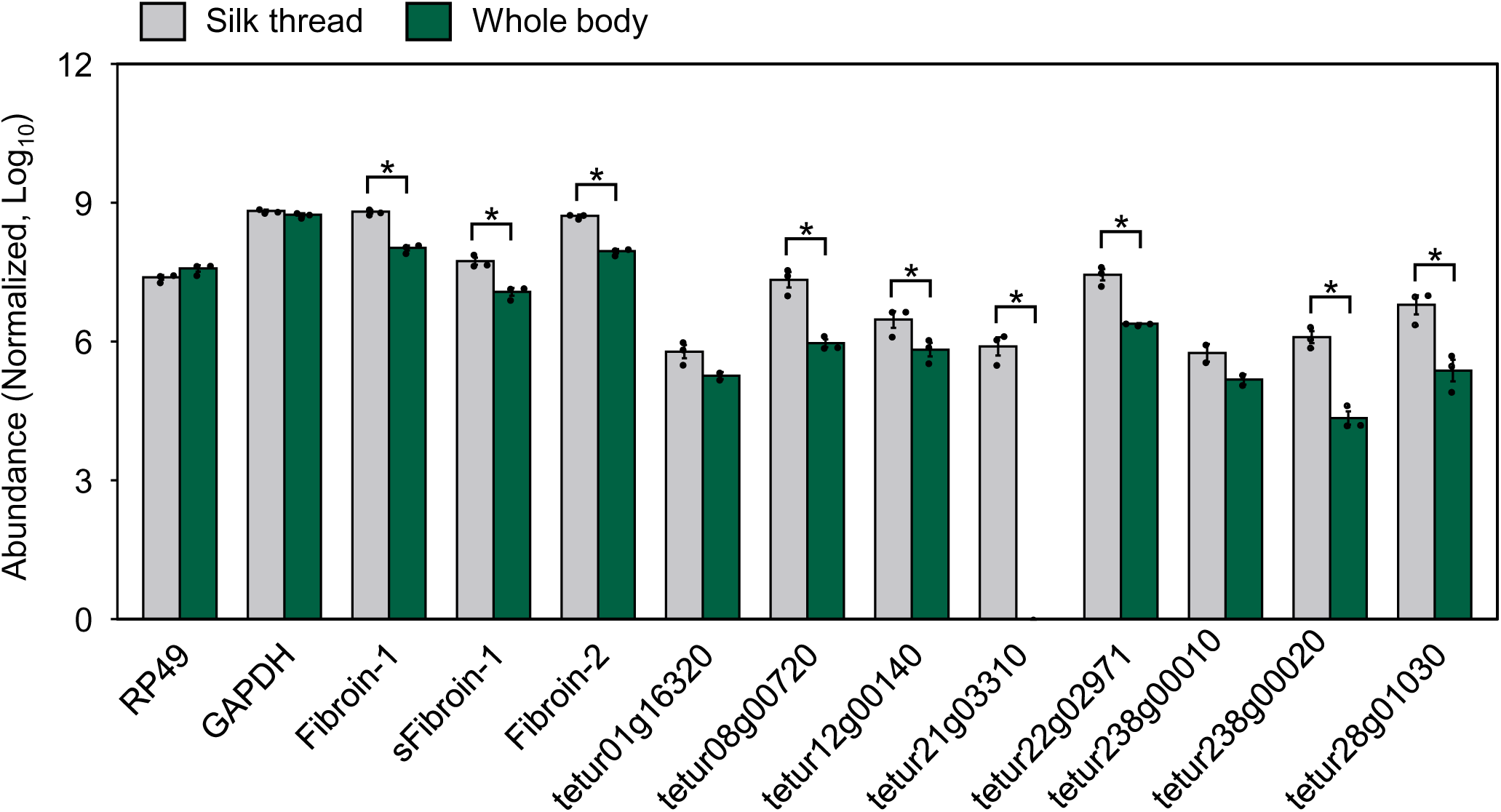
Abundance of proteins detected in silk fiber and the whole body of adult TSSM females. RP49 and GAPDH are the reference proteins. Eight proteins (tetur01g16320 to tetur28g01030) were detected out of 17 candidate silk protein genes.^20^ Data were collected from three independent experimental runs and presented as mean ± SE. In each experimental run, 100-120 and 50-60 individuals were used to collect silk fibers and whole bodies, respectively. * Log2 fold change > 1 and *P* < 0.05. (See also Figure S1)

### *RNAi-mediated functional analysis of* Fibroin-1, sFibroin-1, *and* Fibroin-2 *genes*

We orally administered dsRNA targeting *Fibroin-1* and *sFibroin-1* genes (ds*Fib1*/*sFib1*), *Fibroin-2* gene (ds*Fib2*), or the intergenic region as a negative control (dsNC) to adult TSSM females and evaluated the RNAi-mediated silencing effects. Since *Fibroin-1* and *sFibroin-1* gene sequences have high sequence similarity (72.1%), primers specific for these sequences could not be designed for either real-time RT-qPCR or dsRNA synthesis, so the RNAi-mediated silencing effect could not be distinguished between the two genes by these methods; however, protein expression could be distinguished separately by liquid chromatography-tandem mass spectrometry (LC-MS/MS). Oral administration of ds*Fib1*/*sFib1* or ds*Fib2* reduced the mRNA and protein expression of each target gene compared to oral administration of dsNC (Fig. 4A, B). Compared to oral administration of dsNC, oral administration of a mixture of ds*Fib1*/*sFib1* and ds*Fib2* (ds*Fib*Mix) also reduced the mRNA and protein expression of *Fibroin-1*, *sFibroin-1*, and *Fibroin-2* genes, but did not significantly reduce Fibroin-2 protein expression. Mites treated with ds*Fib1*/*sFib1* showed significantly lower survival (Fig. 4C), fecundity (Fig. 4D), and feeding activity, as assessed by the area of feeding damage formed in leaf discs (Fig. 4E). However, oral administration of ds*Fib2* did not affect these mite performances. A significant positive correlation was found between the fecundity and feeding activity of individual mites (Fig. S2). The duration of a single feeding event was significantly shorter in ds*Fib1*/*sFib1*-treated mites compared to dsNC (Fig. 4F). There was a tendency for the diameter of a single filament to be smaller in mites treated with ds*Fib1*/*sFib1* or ds*Fib2* compared to dsNC (Fig. 4G).

**Figure 4.**
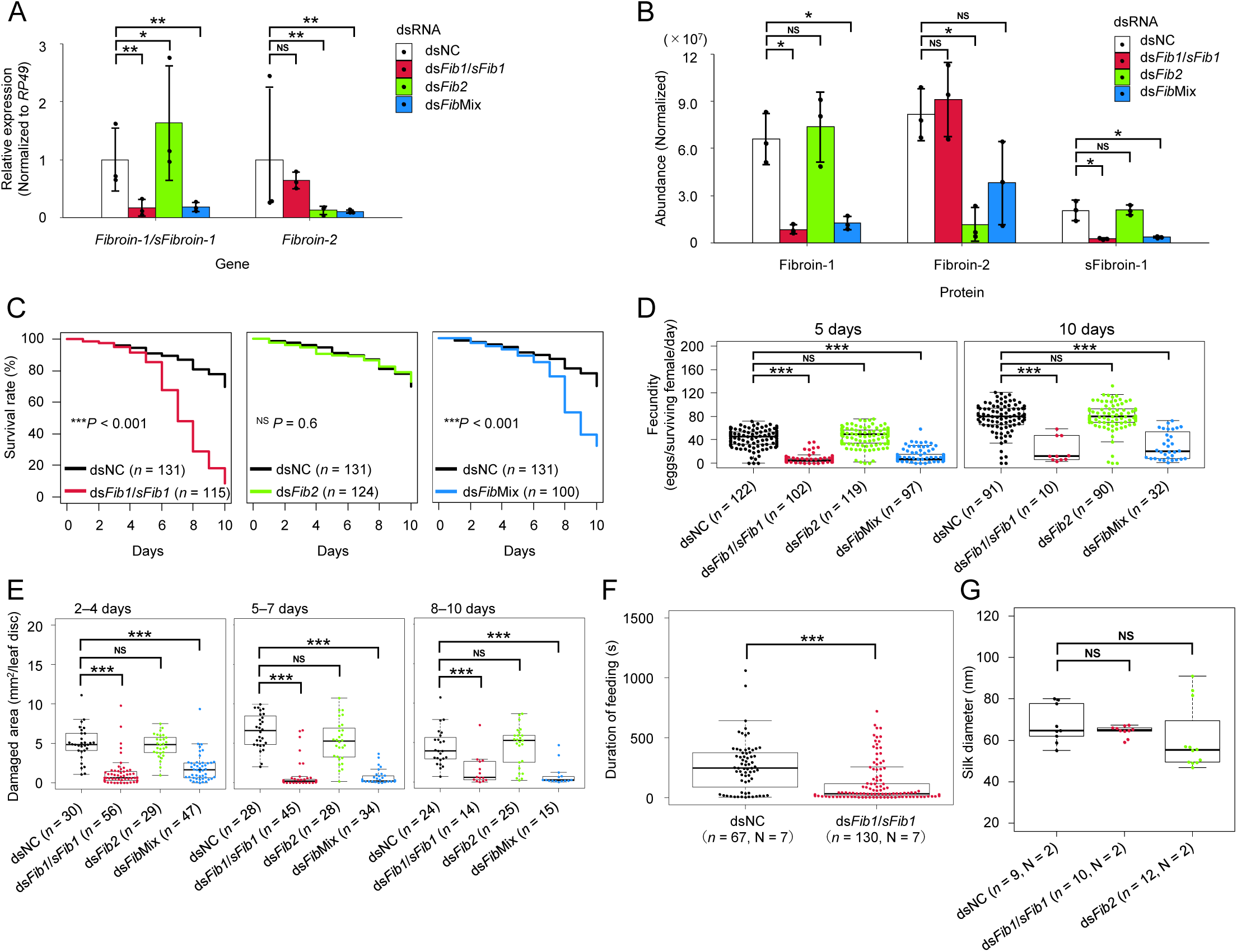
RNAi-mediated silencing of *Fibroin-1*, *sFibroin-1*, and *Fibroin-2* genes affects TSSM performance and silk dimensions. (A) The relative expression levels of *Fibroin-1*/*sFibroin-1* and *Fibroin-2* genes normalized to *RP49* of mites 4 days after RNAi treatment with dsRNA targeting *Fibroin-1*/*sFibroin-1* (ds*Fib1*/*sFib1*), *Fibroin-2* (ds*Fib2*), and the intergenic region as a negative control (dsNC). (B) The abundance of Fibroin-1, Fibroin-2, and sFibroin-1 proteins in mites 4 days after RNAi treatments with ds*Fib1*/*sFib1*, ds*Fib2*, and dsNC. NS, Log2-fold change < 1. *, Log2-fold change > 1 and *P* < 0.05. **, Log2-fold change > 1 and *P* < 0.01. (C) Mite survival for 10 days. (D) Fecundity at 5 and 10 days. (E) Feeding activity every 3 days for 10 days after RNAi treatment with ds*Fib1*/*sFib1*, ds*Fib2*, and dsNC. Survival curves were plotted using the Kaplan-Meier method and compared using the log-rank test (NS, *P* > 0.05; ***, *P* < 0.001). Fecundity and feeding activity were compared using Dunnett’s test with dsNC-treated mites as a negative control (NS, *P* > 0.05; *, *P* < 0.05; **, *P* < 0.01; ***, *P* < 0.001). Data in A-E were collected from three independent experimental runs. (F) Mites’ feeding duration (portion of 1 h) 3 days after RNAi treatment with dsNC and ds*Fib1*/*sFib1* followed by 24 h starvation. Wilcoxon-Mann-Whitney *U* test (***, *P* < 0.001). *n* and *N* indicate the number of feeding events measured and the number of mites used in the experiment, respectively. (G) Dimension of silk filaments from adult mite females 3 days after RNAi treatment with ds*Fib1*/*sFib1*, ds*Fib2*, and dsNC followed by 24 h silk production. Steel’s test (NS, *P* > 0.05). *n* and *N* indicate the number of feeding events measured and the number of mites used in the experiment, respectively. See Figure S2 for correlation analyses.

## Discussion

From the characteristics of repetitive sequences, whole-genome sequencing computationally predicted 17 silk protein genes in TSSM.^20^ Silk proteomics in TSSM identified Fibroin-1 and Fibroin-2, which were not among the 17 candidates, as novel silk proteins,^21^ and these proteins and sFibroin-1, which has 72.1% similarity to Fibroin-1, were also detected in the saliva proteome.^22,23^ In addition, Fibroin-1, sFibroin-1, and Fibroin-2 do not share a high homology with the fibroin protein identified in another spider mite species, *Stigmaeopsis nanjingensis*.^24^ The present study was designed to address the question of whether Fibroin-1, sFibroin-1, and Fibroin-2 function in silk, saliva, or both in TSSM.

The prosomal glands of TSSM consist of a pair of silk glands, a pair of anterior podocephalic glands, a pair of dorsal podocephalic glands, a pair of coxal glands, a pair of tracheal organs, and a single tracheal gland.^25^ Of these, the expression of salivary proteins has only so far been reported in the podocephalic glands, and it is reasonable to assume that these function as salivary glands.^22,23,26–28^ The ISH in the present study revealed the expression of *Fibroin-1*, *sFibroin-1*, and *Fibroin-2* genes in the dorsal podocephalic gland (Fig. 2B–G). This finding is consistent with the results of a previous report on the spatial expression analysis of *Fibroin-1* and *Fibroin-2* genes in TSSM.^22^ Microscopic anatomical analysis using cryo-sectioning and micro-CT techniques showed that the dorsal podocephalic gland was located away from the silk glands (Fig. 2A), and the salivary duct extended to the anterior end of the rostrum without connecting to the silk gland (Fig. 2K, L). These data led us to conclude that, contrary to their names, Fibroin-1, sFibroin-1, and Fibroin-2 are indeed salivary proteins in TSSM.

However, TSSM silk proteomics in the present study showed a high abundance of Fibroin-1, sFibroin-1, and Fibroin-2 (Fig. 3). The detection of Fibroin-1 and Fibroin-2 in silk fibers in TSSM is consistent with the silk proteomics in a previous study.^21^ The expression of these salivary proteins was enriched in the silk fibers compared to the whole mite body, indicating that the salivary proteins detected in the silk fibers were not due to accidental contamination. In fact, of the 95 salivary proteins identified by saliva proteomics,^22^ 66 proteins, including Fibroin-1, sFibroin-1, and Fibroin-2, were detected in the silk proteome in the present study (Fig. S1). Cryo-SEM imaging visualized a beads-on-a-string pattern formed by droplets of fluid on the silk fiber (Fig. 1D, E). Beads-on-a-string and thinning filaments are mainly formed by viscoelastic inertial fluids^29^. They are also known for spider webs^30^ and sticky prey-capturing water-based sugary-proteinaceous fluids, e.g., *Drosera* plants and onychophoran slime^31–34^. This chain-shaped structure is not observed in all TSSM silk fibers. However, the evidence that most salivary proteins were detected in high abundance in the silk fibers (Fig. 3) and that RNAi of the *Fibroin-2* gene tended to produce thin silk filaments (Fig. 4G) suggests that the fluid adhering to the silk fibers is most likely saliva. Saliva may coat the silk fibers and occasionally adheres to the fibers in large amounts, forming the beads-on-a-string pattern. Fibroin-2 has an RQAQA repeat motif^21^ and has properties similar to spider pyriform spidroin and silkworm sericin 4 proteins, which

have high glutamine content.^35–37^ Pyriform spidroin acts as a hardener for the spider’s dragline silk to an object,^38^ and silkworm sericin 4 coats fibroin in the larval silk fiber and provides adhesion.^39^ In addition, secretions derived from the mandibular gland, which produces the majority of the saliva, adhere to silk fibers secreted from the spinnerets in lepidopteran insects.^40^ Furthermore, it is suggested that the chemical compounds contained in the mandibular gland secretions function as a cuticular lipid layer that provides waterproofing to the silk fibers in lepidopteran insects.^41^ These findings suggest that Fibroin-2 may coat TSSM silk fibers and provide adhesive or waterproofing properties.

We found that RNAi-mediated silencing of *Fibroin-1* and *sFibroin-1* genes reduced mite survival, fecundity, and feeding activity (Fig. 4C–E). Mite fecundity and feeding activity in the same individual were positively correlated (Fig. S2), suggesting that suppression of fecundity was due to inhibition of feeding. Most studies of spider mite salivary proteins have focused on elicitor and effector proteins in the context of interactions with plant defense responses.^23,26,27,42–47^ However, it is unlikely that Fibroin-1 and sFibroin-1 have effector functions because SHOT (secreted host-responsive protein of Tetranychidae) family proteins, which have been suggested to have effector functions, showed host plant–dependent expression changes,^23^ but the *Fibroin-1* and *sFibroin-1* gene expression showed no significant differences, at least for feeding on bean, corn, soybean, and tomato plants.^22^ Why did the feeding activity of mites treated with ds*Fib1*/*sFib1* decrease?

Some sucking arthropods use saliva to fix their mouthparts to their hosts. For example, aphids secrete a small amount of gel saliva onto the surface of the host plant to form a flange that provides support for initiating penetration of their stylets into the plant.^48^ As the stylets are inserted through the flange, saliva is transported into the plant tissue to form stylet sheaths that envelop the stylet bundle. The function of stylet sheaths in feeding has been suggested to be to stabilize the stylet and seal the puncture site to prevent leakage of phloem sap into the apoplast.^48–50^ In ticks, the saliva contains Gly-rich flagelliform silk proteins similar to those in spiders that function as cement proteins involved in the blood-sucking behavior.^51,52^ Cryo-SEM visualized a distinct fluid patch remaining at the piercing site on the leaf surface after mite feeding (Fig. 1F, G). This fluid spreads over and wets plant surfaces, also covering anti-adhesive waxes and even embedding them, which lets us deduce the fluid’s contact formation and adhesive properties. A TSSM uses the stylets to inject saliva into the mesophyll cells for pre-oral digestion and to suck the digested contents.^53^ The stylets form a hollow needle consisting of a pair of long movable appendages with a gutter-like structure. Each of these appendages forms a hairpin loop from the stylophore to the anterior end of the rostrum. When the stylophore is retracted, these appendages are pushed out, and the two are interlocked in a longitudinal groove called the rostrum gutter in the middle of the dorsal surface of the rostrum to form a single hollow, needle-shaped channel for feeding.^54^ In the TSSM mouthparts, the stylets and salivary ducts run in parallel to the anterior end of the rostrum (Fig. 2K, L). At its end, the salivary duct joins the stylets,^55^ and saliva presumably flows into the stylets at this point. In the citrus red mite *Panonychus citri* (McGregor.), Tateishi observed the fluid secreted from the anterior end of the rostrum and reported on its adhesion to the leaf surface^56^. When a TSSM placed on a glass plate was observed from the ventral side, it was confirmed that the fluid was probably secreted from the stylets onto the glass surface (Video S1). We hypothesize that the fluid is saliva, which is secreted from the salivary duct through the stylet onto the leaf surface. The saliva may play a role in assisting mite feeding by adhering the anterior end of the rostrum to the leaf surface. The TSSM mouthpart has an organ called the pharyngeal plunger that depressurizes the oral cavity to suck the contents of the mesophyll cells.^57,58^ To achieve this depressurization of the oral cavity using the pharyngeal pump, it may be necessary to make the anterior end of the rostrum adhere to the leaf surface. RNAi-mediated silencing of the *Fibroin-1* and *sFibroin-1* genes reduced the feeding duration in TSSM (Fig. 4F), possibly due to a failure to fix the rostrum to the leaf surface.

The primary structures of Fibroin-1 and sFibroin-1 do not show high similarity to those of the flagelliform silk proteins, but they have silk protein–like properties, including a repetitive pattern of alternating random coil, α-helical and β-strand motifs, and are rich in Val, Asn, and Ser.^21^ These silk-like proteins may provide saliva with sticky properties and support the fixation and stabilizing of the rostrum on the leaf epidermis for TSSM feeding. However, it is unclear why Fibroin-2, which has an RQAQA repeat motif and is thought to coat silk fibers and provide adhesion, is not involved in mite feeding performance (Fig. 4F). On the other hand, although Fibroin-1 and sFibroin-1 are also frequently detected in silk fibers, it is also unclear why RNAi of these genes did not affect their dimensions (Fig. 4G). In addition, Fibroin-1 and Fibroin-2 were detected in the TSSM fecal proteome.^59^ TSSM feces are derived from digestive cells that are responsible for intracellular digestion while floating in the midgut.^60^ Since the expression of *Fibroin-1* and *Fibroin-2* genes was not detected in tissues other than the dorsal podocephalic gland (Fig. 2B–J), it is possible that they are secreted as saliva and then transferred to the digestive system, where they are taken up by digestive cells. The function of these proteins in the digestive process is unknown.

Eight of the 17 predicted silk protein candidates were also detected in the TSSM silk proteomics in the present study (Fig. 3), but these proteins were not detected by Arakawa et al. in the previous silk proteomics.^21^ The abundance of these eight proteins was lower than that of Fibroin-1, sFibroin-1, and Fibroin-2. Whereas Arakawa et al. collected TSSM silk fibers produced on leaves, we placed starved adult females on empty Petri dishes and collected silk fibers suspended in the air. Our collection method had less contamination from molted shells and feces than the method used by Arakawa et al., which may have contributed to the detection of these candidate silk proteins. We suggest that these detected proteins have the potential to be functional silk proteins of TSSM. A limitation of the present study is that there are no data to explain the detailed mechanism of saliva adhesion to the silk fibers in TSSM. It is possible that the liquid silk secreted from the spinnerets is spun, and then the fibers are submerged in the saliva secreted from the stylets before contacting the leaf surface because the anterior end of the rostrum, where the stylet exits, is adjacent to the spinnerets. However, this is a hypothesis. It is unknown whether saliva is thinly and uniformly coated on the silk fibers outside of the beads-on-a-string pattern region. To clarify this, molecular labeling of saliva and silk using TSSM transformation^61^ would be effective. Interestingly, silk fibers frequently carried attached epicuticular plant wax crystals (Fig. 1F), which must have adhered to sticky (saliva-covered) sites of the silk and detached upon silk tension. This observation supports our hypotheses above.

In summary, Fibroin-1, sFibroin-1, and Fibroin-2 are salivary proteins that are produced in the salivary glands, are secreted from the stylets through the salivary duct, and adhere to silk fibers in TSSM. The functions of Fibroin-1 and sFibroin-1 may be to adhere the mouthpart to the leaf surface to stabilize the stylets when sucking plant cell contents. In addition, Fibroin-2 may act as a coating agent for silk fibers. The mechanism by which TSSM saliva coats silk fibers, and the physical properties of the saliva itself and the saliva-coated silk fibers must be experimentally confirmed and require further study.

## Materials and Methods

### Mites

We used two populations of TSSM. One population was collected in London, Ontario, Canada, and was previously used for whole-genome sequencing.^20^ This population was maintained on kidney bean *Phaseolus vulgaris* L. ’Hatsumidori #2’ plants in our laboratory at an air temperature of 25 °C and a light period of 16 h each day. The other population was collected from a strawberry field in Weixdorf (Saxony, Germany)^62–64^, maintained on broad bean *Vicia faba* L. ’Witkiem’, and used for cryo-SEM observations. Therefore, mites were transferred to and established on 5-leaf-stage plants of *Fragaria chiloensis* (L.) Mill. and *Solanum lycopersion* L. ’Harzfeuer’. The London population was used for all other experiments.

### SEM and cryo-SEM

To visualize TSSM mouthparts, silk fibers, and saliva, we used SEM combined with a VHX-D500 microscope (Keyence Corp., Osaka, Japan) and the cryo-SEM FE-SEM Supra 40VP-31-79 microscope (Carl Zeiss SMT, Oberkochen, Germany) equipped with an Emitech K250X cryo-preparation unit (Quorum Technologies Ltd., Ashford, Kent, UK). For cryo-SEM, cut pieces of fresh, just fully developed, mite-infested leaves were mounted on metal holders with Tissue-Tek O.C.T. Compound (Sakura Fine Technical Co., Ltd., Tokyo, Japan), and then frozen in the cryo-preparation unit at -140 °C, sputter-coated with a 6-nm thick layer of platinum, and examined in the frozen state inside the cryo-SEM unit at -90 °C and 5 kV acceleration voltage.

### AFM

TSSM silk fibers on a silicon wafer (4136SC; Alliance Biosystems, Osaka, Japan) were observed using an AFM (SPM-9600; Shimadzu, Kyoto, Japan). A single mite 3 days after RNAi treatment was allowed to move around the silicon wafer at room temperature for 24 h. After 24 h, the mite was removed, and silk fibers remaining on the silicon wafer were observed. The dimensions of individual filaments of silk fiber were analyzed from images obtained in the tapping mode.

### ISH by RNAscope

Approximately 200 randomly aged female TSSM were collected in 1.5-mL centrifuge tubes using an air pump-based mite collection system.^65,66^ Mites were fixed with 100% MeOH for 12 h. Then, the solution was replaced with 15% and 30% sucrose solutions in sequence. The samples were embedded in Tissue-Tek O.C.T. Compound (Sakura Finetek Japan Co., Ltd., Tokyo, Japan), frozen at −100 °C, and stored at −80 °C for 12 h. Frozen, 20-μm thick sections were cut on glass slides and incubated for 12 h at 37 °C. Sections were stored at −80 °C until used for ISH. *Fibroin-1*, *sFibroin-1*, and *Fibroin-2* expression sites were analyzed using an RNAscope (Advanced Cell Diagnostics Inc., Hayward, CA),^67,68^ an ISH technique that detects mRNA with high sensitivity and specificity. Probes targeting *Fibroin-1* (456–3960 bp, LOC107361698), *sFibroin-1* (270–3331 bp, LOC107361899), and *Fibroin-2* (131–1280 bp, LOC107368919) were designed by Advanced Cell Diagnostics (Newark, CA, USA). The *Fibroin-1* probe was used in the C1 channel, and *sFibroin-1* and *Fibroin-2* in the C2 channels. The RNAscope 3-plex Negative Control Probe was used as a negative control. Sections were fixed with 4% paraformaldehyde for 15 min at 4 °C, washed with 1×PBS, and then dehydrated by immersion in 50%, 70%, and 100% ethanol for 5 min each. Hybridization was performed according to the protocol of the RNAscope Multiplex Fluorescent Reagent Kit Ver.2. Sections were treated with hydrogen peroxide (supplied with the kit) for 10 min at room temperature and washed with water. To enhance signal intensity, target retrieval pretreatment was performed according to the manufacturer’s protocol. The sections were treated with Protease Ⅳ (supplied with the kit) at room temperature for 30 min and washed with 1× PBS. Probes were treated at 40 °C overnight and washed with 1× wash buffer (twice for 2 min). The sections were treated with AMP1, followed by AMP2 at 40 °C for 30 min and AMP3 at 40 °C for 15 min (AMP1, 2, and 3 were supplied with the kit). Opal 520 (Akoya Biosciences, Marlborough, MA, USA) was used to detect the C1 channel probe (green), and Opal 570 (Akoya Biosciences) was used to detect the C2 channel probe (red). After treatment with horseradish peroxidase (HRP)-C1 or HRP-C2 (supplied with the kit) for 15 min at 40 °C, the sections were treated with 1000-fold diluted Opal 520 or Opal 570 for 30 min at 40 °C. To stop the HRP reaction, the sections were treated with HRP blocker (supplied with the kit) for 15 min at 40 °C. The sections were mounted with deionized water and coverslips and observed under a confocal laser scanning microscope FV10i (Olympus, Tokyo, Japan).

### Micro-CT

Approximately 200 randomly aged female TSSM were collected into 1.5-mL centrifuge tubes using an air pump-based mite collection system.^65,66^ Mites were washed with PTw (PBS with 0.1% Tween-20) (three times for 5 min) and then were fixed with slight shaking in 4% paraformaldehyde for 12 h at room temperature. For dehydration, mites were washed with 1xPBS, then 50% and 70% ethanol were added sequentially, and the samples were left at 25 °C for 10 min. Ethanol with 0.3 M I_2_ and 1% phosphotungstic acid were added, and the samples were stained at 25 °C for more than 12 h. The well-stained mites were placed in small plastic bags containing 70% ethanol (5 mites per bag), and the positions of the mites were fixed with fibers from wipes (Kimwipe; Kimberly-Clark, Irving, TX) to prevent mites from overlapping. Each bag was heat-sealed to prevent air from entering. The mites in the bags were aligned vertically on a stand, and images were taken using a three-dimensional X-ray microscope (Xradia 510 Versa; Carl Zeiss AG, Jena, Germany; 50 kV, 79 μA).

### LC-MS/MS

Approximately 50 mites that had been RNAi-treated 4 days previously were collected in 1.5-mL centrifuge tubes using an air pump–based mite collection system^65,66^ and used for expression analysis. Silk fibers were collected as follows: approximately 100 mites of random age that had been starved for 1 day were placed on a plastic Petri dish (diameter 35 mm) for 1 day, and the silk produced in the air was rolled up with a plastic homogenizer pestle. For protein extraction, protein lysis buffer containing 10 mM Tris-HCl (pH 9.0) and 8 M urea was added to the tube containing the mites, and protein lysis buffer containing 10 mM Tris-HCl (pH 9.0) and 6 M guanidine was added to the tube containing the silk fibers. The mites or silk fibers were ground using a plastic homogenizer pestle and an ultrasonic cleaner (AS38A; As one, Osaka, Japan). The tube containing the mites was centrifuged at 12,000 rpm for 10 min at 4 °C (Eppendorf model 5424 R centrifuge; Eppendorf AG, Hamburg, Germany) to remove cell debris, and the supernatant was used in the following steps. The concentration of the extracted proteins was measured using a Qubit 4 fluorometer (Thermo Fisher Scientific, Waltham, MA, USA). The extracted proteins were reduced with 10 mM DTT (Invitrogen, Waltham, MA, USA) for 30 min and then alkylated with 50 mM IAM (Fujifilm Wako Pure Chemical, Osaka, Japan) for 20 min in the shade. They were then diluted with 50 mM NH_4_HCO_3_ (Fujifilm Wako Pure Chemical, Osaka, Japan) to a quarter urea concentration, followed by digestion into peptides with trypsin (Promega K.K., Tokyo, Japan) (trypsin:protein = 1:50) at 25 °C for 12 h. To stop the trypsin reaction, 2% TFA was added in the same volume as the sample solution and centrifuged at 13,000 rpm, 4 °C for 10 min. Solution A (water:ACN:TFA = 200:800:1) was added to 200 µL of peptide desalting column (GL-Tip SDB; GL Sciences Inc., Tokyo, Japan) and centrifuged at 3400 rpm, 4 °C for 3 min, followed by the addition of solution B (water:ACN:TFA = 950:50:1) under the same conditions. The sample was then dropped into the column, washed with solution B, and the digested protein eluted with solution A. Samples were concentrated in a vacuum evaporator and stored at −80 °C until applied to the LC-MS/MS.

Shotgun analysis was performed on an Orbitrap Exploris 480 (Thermo Fisher Scientific). The raw mass spectrometry data were deposited as JPST003501 for jPOSTrepo (https://repository.jpostdb.org/)^69^ and as PXD059809 for ProteomeXchange (https://www.proteomexchange.org/).^70^ The MS data were analyzed using Proteome Discoverer v.2.5.0.400 software (Thermo Fisher Scientific). The protein database was TSSM annotation (v.20190125) from the ORCAE database.

### dsRNA synthesis and delivery

The nucleotide sequences of *Fibroin-1* (tetur07g00160), *sFibroin-1* (tetur07g01660), and *Fibroin-2* (tetur29g01360) genes were obtained from the ORCAE database (https://bioinformatics.psb.ugent.be/orcae/overview/Tetur). Total RNA was extracted from the frozen female mites using NucleoSpin RNA Plus XS (Macherey-Nagel GmbH & Co. KG, Düren, Germany) according to the manufacturer’s protocol. The concentration of the extracted RNA was measured using a spectrophotometer (NanoPhotometer N60; Implen, Munich, Germany). cDNA was reverse-transcribed from the extracted total RNA using a SuperScript II cDNA Synthesis Kit (Thermo Fisher Scientific) and stored at −30 °C. Genomic DNA (gDNA) was extracted using a NucleoSpin Tissue Extraction Kit (Macherey-Nagel) and stored at −30 °C. Using cDNA as a template, a 601-bp fragment of both *Fibroin-1* and *sFibroin-1* and a 648-bp fragment of *Fibroin-2* (Fig. S3) were PCR-amplified using KOD DNA polymerase (Toyobo, Osaka, Japan). A 382-bp intergenic fragment (non-coding [NC], genomic coordinates: scaffold 12, position 1690614–1690995; Fig. S3)^71^ was PCR-amplified in the same manner using gDNA as a template. Primers to amplify the DNA fragments of *Fibroin-1*, *Fibroin-2*, and the NC fragment are shown in Table S1. Amplified DNA fragments were purified using NucleoSpin Gel and PCR Clean-Up Kit (Macherey-Nagel). RNA fragments were transcribed from each DNA template using an *in vitro* Transcription T7 Kit (Takara Bio, Kusatsu, Japan) in 1.5-mL centrifuge tubes. After treatment with DNase Ⅰ (Takara Bio) for 30 min, the RNA fragments were denatured at 95 °C for 5 min, followed by a slow cool-down to 25 °C to facilitate the formation of dsRNA. The dsRNA fragments (ds*Fib1*/*sFib1*, ds*Fib2*, and dsNC) were purified by phenol-chloroform extraction and ethanol precipitation.

A paraffin wax film (Parafilm M; Bemis, Neenah, WI, USA) folded in four was placed on the bottom of a plastic Petri dish, and the four corners were crimped. Ten microliters of dsRNA solution (1 µg μL^−1^; 0.1% (v/v) Tween 20; 0.1% blue dye) were pipetted onto a piece of Kimwipe on the parafilm. Fifty newly molted adult females were placed with their ventral side in contact with the Kimwipe and incubated at 25 °C for 2 h. The digestive system turned blue due to the blue dye (Brilliant Blue FCF; Fujifilm Wako Pure Chemical, Osaka, Japan) contained in the dsRNA solution. Individuals that turned blue were confirmed to have delivered the dsRNA solution, selected under a stereomicroscope, and transferred onto the leaves. They were used in each experiment immediately after RNAi treatment or after 4 days of rearing on the leaves.

### Real-time RT-qPCR

Four days after RNAi treatment, total RNA was extracted from 50 mites in each of three independent experimental runs using NucleoSpin RNA Plus XS. cDNA was reverse-transcribed from the extracted total RNA using the High Capacity cDNA Reverse Transcription Kit (Thermo Fisher Scientific). qRT-PCR reactions were performed in three technical replicates using Power SYBR Green Master Mix (Thermo Fisher Scientific) on an ABI StepOnePlus Real-Time PCR System (Thermo Fisher Scientific). The reference gene *RP49* (tetur18g03590), which encodes a ribosomal protein, was used as an internal control. Primer sequences and amplification efficiencies for the reference gene and target gene are shown in Table S2. The cycle threshold value of each independent experimental run was calculated from the average of 3 technical replicates, respectively. The expression value for each target gene was normalized to the reference gene. The normalized relative quantity (NRQ) was calculated as follows: NRQ = (1+E_R_)^CtR^/(1+E_T_)^CtT^, where E_R_ and E_T_ are the amplification efficiencies for reference and target genes, respectively, and CtR and CtT are the cycle threshold value of reference and target genes, respectively.

### Mite performance assay

Immediately after RNAi treatment, mites were transferred onto fresh kidney bean–leaf discs (1 mite/disc). Mite survival and fecundity were observed daily for 10 days. Leaf discs were replaced with fresh ones every three days, and the area of mite feeding damage formed in each three-day period was quantified by image analysis^65^ as feeding activity. The feeding duration of TSSM was measured according to a previously described method^72^. Three days after RNAi treatment, mites were fasted for 24 h in a plastic Petri dish. A single mite was placed on a kidney bean–leaf disc (8 mm diameter) and captured on video for 1 h using a stereomicroscope (S8APO; Leica Microsystems GmbH, Wetzlar, Germany) equipped with a digital camera (EOS Kiss X7; Canon Inc., Tokyo, Japan).

## Resource availability

### Lead contact

Further information and requests for resources should be directed to and will be fulfilled by the lead contact, Takeshi Suzuki.

### Materials availability

This study did not generate new unique reagents.

### Data and code availability

- Proteomics data have been deposited at jPOSTrepo (accession number: JPST003501) and ProteomeXchange (accession number: PXD059809), and are publicly available as of the date of publication.
- This paper does not report the original code.
- Any additional information required to reanalyze the data reported in this paper is available from the lead contact upon request.

## Acknowledgments

The authors would like to thank Dr. K. Arakawa of Keio University for their advice on LC-MS/MS. We also thank Dr. Y. Kamei of the National Institute for Basic Biology and Dr. S. Shimano of Hosei University for their assistance with micro-CT imaging. DV appreciates the access to cryo-SEM provided by Prof. Dr. Christoph Neinhuis, the Chair of Botany at the faculty of Biology of TU Dresden, Germany. This work was supported in part by JSPS KAKENHI (grant numbers 18H02203, 21H02193, and 24K21256) awarded to T.S., and in part by the Cabinet Office, Government of Japan Cross-ministerial Moonshot Agriculture, Forestry and Fisheries Research and Development Program, “Technologies for Smart Bio-industry and Agriculture” (funded by the Bio-oriented Technology Research Advancement Institution) (JPJ009237) awarded to T.S..

## Author contributions

Conceptualization, Y.A., N.T., and T.S.; methodology, Y.A., N.T., S.K., H.M., D.V., and T.S.; investigation, Y.A., N.T., and D.V.; software, Y.A., N.T., and T.S.; formal analysis, Y.A. and N.T.; visualization, Y.A., N.T., and D.V.; writing – original draft, Y.A., N.T., D.V., and T.S.; writing – review and editing, Y.A., N.T., D.V., and T.S.; supervision, T.S.; funding acquisition, T.S.

## Declaration of interests

The authors declare no competing interests.

## Supplemental information

Document S1: Figures S1–S3 and Tables S1–S3

Video S1. Fluid secretion observed from the ventral side of a TSSM.

---

**Figure S1. Heat map of salivary and silk protein abundance in silk fibers and whole body, for data used in Figure 3**

The numbers after the sample name (Silk and Mites) indicate biological replicates. Of the 95 proteins previously identified in the salivary proteome,^22^ 66 proteins (shown in blue on the left) were detected in at least one silk sample. Of the 17 silk protein candidates predicted from the genome,^20^ 8 proteins (shown in green) were detected in at least one silk sample. The heat map colors indicate the abundance of each protein: high expression is red, low expression is orange, and undetected is gray. This heat map was generated using the R package Superheat (https://rlbarter.github.io/superheat/).

**Figure S2. Correlation between fecundity and feeding activity in the data for RNAi-treated mites used in Figure 4**

Spearman’s correlation coefficients (*r*) were calculated for the relationship between mite fecundity and area damaged by mite feeding on kidney-bean leaf discs at 2–4, 5–7, and 8–10 days after RNAi treatment with dsNC, ds*Fib1*/*sFib1*, ds*Fib2*, or a mixture of ds*Fib1*/*sFib1* and ds*Fib2*(ds*Fib*Mix). Data were collected from three independent experimental runs. Parameters of the correlation and linear regression analyses are shown in Table S3.

**Figure S3. Gene structures of RNAi targets**

Gene IDs for *Fibroin-1*, *sFibroin-1*, *Fibroin-2* genes are tetur07g00160, tetur29g01360, and tetur07g01660, respectively. Red boxes indicate the regions used for the synthesis of dsRNA targeting *Fibroin-1*/*sFibroin-1* (ds*Fib1*/*sFib1*, 601 bp), *Fibroin-2* (ds*Fib2*, 648 bp), and the intergenic region in scaffold 12 of the TSSM genome as negative control (dsNC, 382 bp^62^).

**Video S1. Fluid secretion observed from the ventral side of a TSSM**

The stylet appears and taps the glass plate from the second 4 in the video. The fluid secretion remaining on the glass plate and the silk fiber extending from the fluid can be seen between seconds 17 and 22.

## Notes

https://figshare.com/articles/media/Salivary_proteins_in_spider_mites_dual_roles_in_feeding_and_silk_fiber_coating/28374404?file=52211765

### Competing Interest Statement

The authors have declared no competing interest.

