## Supplementary material for "Salivary proteins in spider mites: dual roles in feeding and silk fiber coating": Figures S1-S3 and Tables S1-S3

Figure S1

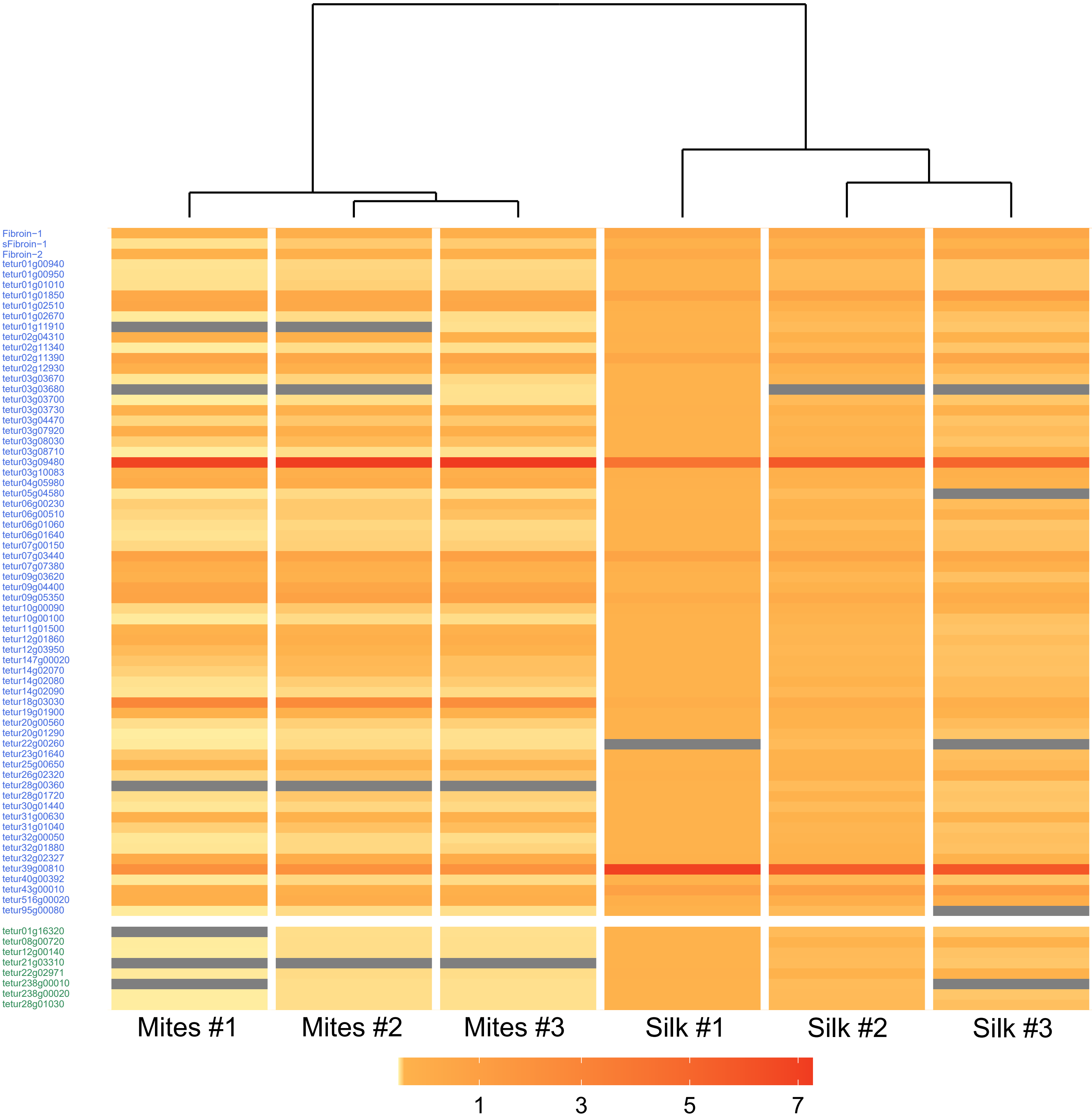
Fibroin−1
sFibroin−1
Fibroin−2
tetur01g00940
tetur01g00950
tetur01g01010
tetur01g01850
tetur01g02510
tetur01g02670
tetur01g11910
tetur02g04310
tetur02g11340
tetur02g11390
tetur02g12930
tetur03g03670
tetur03g03680
tetur03g03700
tetur03g03730
tetur03g04470
tetur03g07920
tetur03g08030
tetur03g08710
tetur03g09480
tetur03g10083
tetur04g05980
tetur05g04580
tetur06g00230
tetur06g00510
tetur06g01060
tetur06g01640
tetur07g00150
tetur07g03440
tetur07g07380
tetur09g03620
tetur09g04400
tetur09g05350
tetur10g00090
tetur10g00100
tetur11g01500
tetur12g01860
tetur12g03950
tetur147g00020
tetur14g02070
tetur14g02080
tetur14g02090
tetur18g03030
tetur19g01900
tetur20g00560
tetur20g01290
tetur22g00260
tetur23g01640
tetur25g00650
tetur26g02320
tetur28g00360
tetur28g01720
tetur30g01440
tetur31g00630
tetur31g01040
tetur32g00050
tetur32g01880
tetur32g02327
tetur39g00810
tetur40g00392
tetur43g00010
tetur516g00020
tetur95g00080
tetur01g16320
tetur08g00720
tetur12g00140
tetur21g03310
tetur22g02971
tetur238g00010
tetur238g00020
tetur28g01030
Mites #1
Mites #2
Mites #3
Silk #1
Silk #2
Silk #3
1
3
5
7

### Figure S2

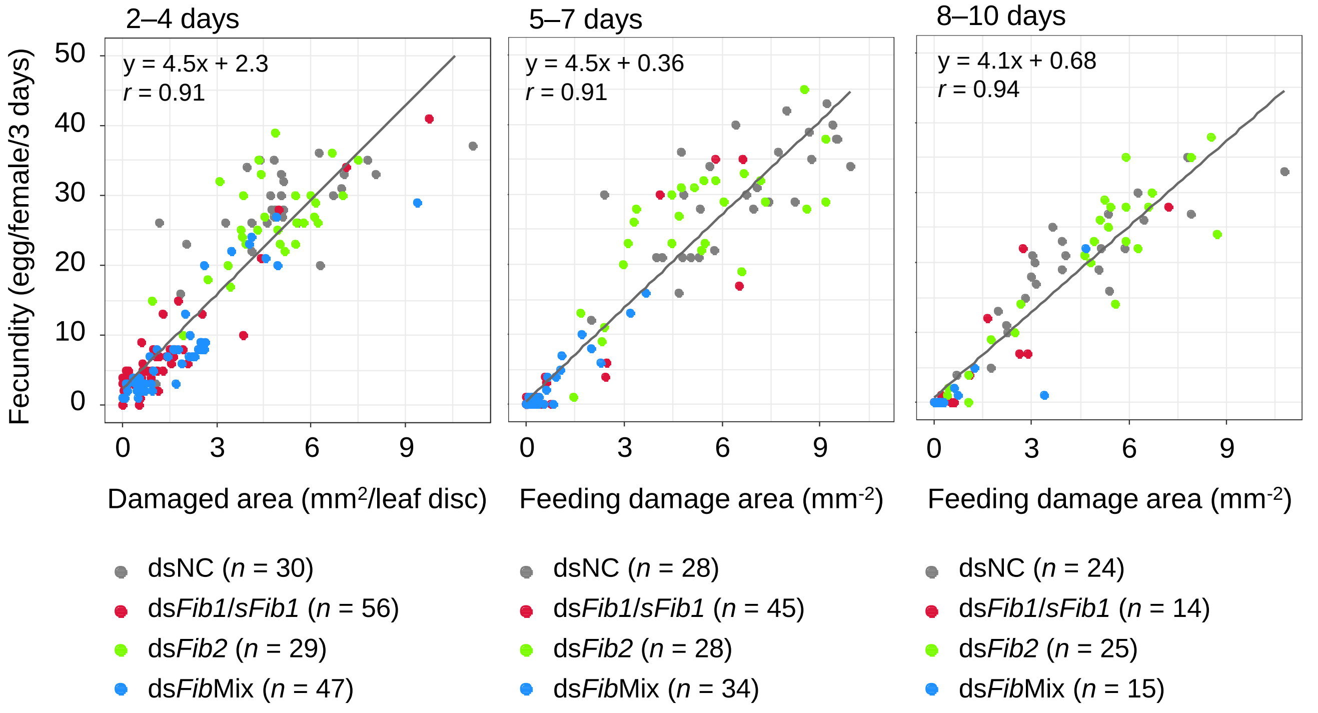

2–4 days
5–7 days
8–10 days
y = 4.5x + 2.3
r = 0.91
y = 4.5x + 0.36
r = 0.91
y = 4.1x + 0.68
r = 0.94
Fecundity (egg/female/3 days)
0
10
20
30
40
50
0
3
6
9
Damaged area (mm2/leaf disc)
Feeding damage area (mm-2)
Feeding damage area (mm-2)
dsNC (n = 30)
dsFib1/sFib1 (n = 56)
dsFib2 (n = 29)
dsFibMix (n = 47)
dsNC (n = 28)
dsFib1/sFib1 (n = 45)
dsFib2 (n = 28)
dsFibMix (n = 34)
dsNC (n = 24)
dsFib1/sFib1 (n = 14)
dsFib2 (n = 25)
dsFibMix (n = 15)

Figure S3

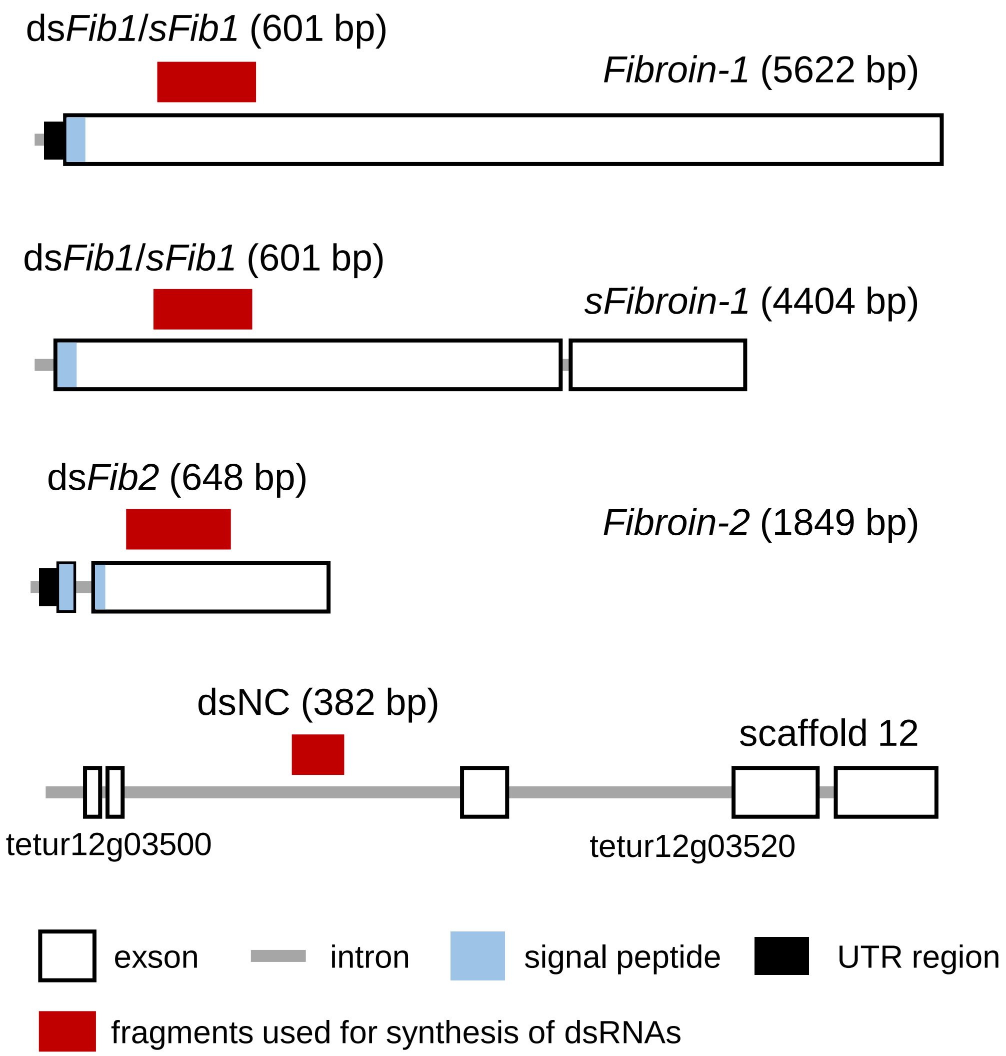
dsFib1/sFib1 (601 bp)
Fibroin-1 (5622 bp)
dsFib1/sFib1 (601 bp)
sFibroin-1 (4404 bp)
dsFib2 (648 bp)
Fibroin-2 (1849 bp)
dsNC (382 bp)
scaffold 12
tetur12g03500
tetur12g03520
exson
intron
signal peptide
UTR region
fragments used for synthesis of dsRNAs

**Table S1. Primers for dsRNA synthesis**

| Target | 5'–3' Forward primer sequence | 5'–3' Reverse primer sequence | Fragment size (bp) |
|---|---|---|---|
| *Fibroin-1/sFibroin-1* | TAATACGACTCACTATAGGGAGCAGCTTGGTATCGTCTGA | TAATACGACTCACTATAGGGTTCATCAGCAACCCCAGTC | 601 |
| *Fibroin-2* | TAATACGACTCACTATAGGGTTGGTTTCGGATCTGGCTCT | TAATACGACTCACTATAGGGCTTGACGTTGGATCTGGGC | 648 |
| NC | TAATACGACTCACTATAGGGCGACCCCATCAGGCTATTGA | TAATACGACTCACTATAGGGGCCCTCTCCTGGTTGTAAACTT | 382 |

Underlined regions indicate the T7 promoter sequence. NC, intergenic region as a negative control.

**Table S2. Primers for real-time RT-qPCR**

| Target | 5'–3' Forward primer sequence | 5'–3' Reverse primer sequence | Primer efficiency (%) |
|---|---|---|---|
| *Fibroin-1/sFibroin-1* | ACAAATTCGCCCGTTCAC | TGAGTTTCGGATGTCAGAGGA | 103.8 |
| *Fibroin-2* | CCTCAAGCTCCCGAGGTTTC | CCTTGAACTCCACCAGCAC | 103.8 |
| *RP49* | TGGTTACGGATCAGCCAAAG | GAATGAGCGATTTCACCACA | 99.1 |

**Table S3. Parameters of correlation and linear regression analyses shown in Figure S2**

| | | Correlation analysis | | Linear regression analysis | | |
|---|---|---|---|---|---|---|
| Days | Treatment | Spearman's Rho | *P* value | Slope | Intercept | *P* value |
| 2–4 | All | **0.91** | <2.2e-16 | 4.5 | 2.3 | <2.2e-16 |
| | dsNC | 0.61 | 3.8e-04 | 2.4 | 16 | 6.7e-05 |
| | ds*Fib1*/*sFib1* | **0.83** | 3.1e-15 | 4.2 | 1.8 | <2.2e-16 |
| | ds*Fib2* | 0.54 | 2.5e-03 | 2.7 | 13 | 2.7e-04 |
| | ds*Fib*Mix | **0.90** | <2.2e-16 | 3.9 | 1.1 | <2.2e-16 |
| 5–7 | All | **0.91** | <2.2e-16 | 4.5 | 0.36 | <2.2e-16 |
| | dsNC | **0.71** | 2.1e-05 | 2.8 | 12 | 8.5e-06 |
| | ds*Fib1*/*sFib1* | 0.62 | 5.2e-06 | 4.8 | -0.67 | <2.2e-16 |
| | ds*Fib2* | **0.77** | 2.0e-06 | 3.7 | 6.5 | 4.2e-08 |
| | ds*Fib*Mix | **0.70** | 4.1e-06 | 4.3 | -0.56 | <2.2e-16 |
| 8–10 | All | **0.94** | <2.2e-16 | 4.1 | 0.68 | <2.2e-16 |
| | dsNC | **0.86** | 7.0e-08 | 2.9 | 7.2 | 1.1e-07 |
| | ds*Fib1*/*sFib1* | **0.83** | 2.1e-04 | 4.2 | −0.35 | 1.2e-05 |
| | ds*Fib2* | **0.84** | 1.8e-07 | 4.2 | 0.62 | 1.5e-10 |
| | ds*Fib*Mix | **0.82** | 2.1e-04 | 3.4 | −0.73 | 1.9e-04 |

Bold numbers indicate Spearman's Rho > 0.70.
